# Gene network switching provides a mechanistic basis for time-to-space translation in insect embryonic patterning

**DOI:** 10.1101/2025.09.10.675428

**Authors:** Mahla Ahmadi, Heike Rudolf, Christine Mau, Jimena Garcia-Guillen, Ezzat El-Sherif

## Abstract

The French Flag model has long served as the prevailing framework for explaining how morphogen gradients generate spatial domains during embryonic development. More recently, however, evidence has shown that many tissues instead establish patterns by translating the sequential activation of genes (organized into genetic cascades) into spatial domains. This translation is thought to occur through modulation of the speed or timing of cascade progression, but the mechanisms underlying such temporal control remain unclear. Two models have been proposed: the *general kinetic modulation* model, in which morphogens influence global kinetic factors such as transcription and decay rates, and the *gene regulatory network (GRN) switching* model, in which morphogens reconfigure regulatory interactions so that genes initially function within a genetic cascade driving sequential activation, but are later integrated into a stabilizing network that locks their expression into mutually reinforcing domains. This transition is hypothesized to occur through a shift from a dynamic GRN (a genetic cascade that drives sequential activations) to a static GRN (a stabilizing network that maintains gene expression domains). Using gap genes in Tribolium castaneum as a model, we combined HCR in situ hybridization, parental RNA interference, and computational modeling to test these hypotheses. We show that gap genes initially act in a genetic cascade producing sequential activations, followed by a morphogen-dependent reconfiguration that stabilizes spatial domains. In particular, we identify the Mlpt-Svb complex as a key stabilizing factor that maintains *svb* expression anteriorly after its initial activation in the posterior. Computational simulations reproduce experimental phenotypes and support GRN switching as the underlying mechanism. Together, these findings demonstrate how morphogen-driven rewiring of network interactions converts temporal cascades into stable spatial patterns, providing a mechanistic basis for robust anterior– posterior patterning in insect embryos and beyond.

## Introduction

Embryogenesis is an intricate process wherein a single fertilized egg undergoes extensive cell divisions and differentiations to form a complete organism. A pivotal step in this process is the establishment of cell identities, where individual cells adopt specific functions and occupy a precise location within the embryo. Understanding Gene Regulatory Networks (GRNs) that regulate these cell identities is vital for comprehending how organisms develop their body plans and structures (1,2). These networks involve the activation and interaction of specific genes that define cell fate in response to positional information, ensuring proper tissue and organ formation during development.

One widely used model to explain body patterning during embryogenesis is the French flag model (3). According to this model, cells read the concentration of morphogens, and depending on whether the concentration crosses certain thresholds, they activate different genes. This creates sharp boundaries between gene expression domains, helping cells know their position and fate within the developing embryo. While this model has been foundational, it assumes that morphogen gradients are stable and produce precise thresholds. Recent studies, however, show that these gradients are often noisy and dynamic (4,5), challenging the idea that fixed concentration thresholds alone can reliably direct consistent gene expression during development.

The speed regulation model (6,7) offers an alternative to the French flag model by focusing on the timing of gene activation rather than fixed morphogen concentration thresholds. In this model, each cell undergoes successive activations of genes, with a morphogen (referred to as the ‘speed regulator’) controlling the rate at which these activations occur. This results in a gradual transition of gene expression rather than sharp, static boundaries. The model translates temporal rhythms in gene activities, where genes are activated sequentially (or periodically) over time, into spatial patterning. In this scheme, cells exposed to different morphogen concentrations activate genes at different speeds, creating a more dynamic and flexible system, where positional information is encoded in the speed of gene activation, not in fixed concentration thresholds. The speed regulation model operates in two complementary forms. The first, the gradient-based form, uses a morphogen gradient to modulate the speed of gene activation across cells: high morphogen concentrations trigger rapid gene activation, while lower concentrations slow it down. This creates waves of gene expression that propagate from regions of high morphogen concentration to lower ones, eventually stabilizing spatial expression domains as the gradient declines. In the second, the wavefront-based form, temporal gene activations proceed autonomously across the tissue, often in the form of oscillations or sequential cascades. A moving boundary of the speed regulator, or “wavefront,” then sweeps across the tissue and progressively freezes cells in their current state. Cells ahead of the wavefront continue cycling through gene activations, while those behind it are locked into a specific fate, converting temporal dynamics into a spatially ordered pattern.

This type of temporal modulation has been implicated in patterning various embryonic tissues, like the segmentation of AP axis in arthropods and vertebrates (7–16), Hox gene regulation in vertebrates (7,17,18) and gap gene regulation in insects (6,19,20), and the patterning of ventral neural tube (21,22) and limb bud (23,24) in vertebrates. However, while the speed regulation model explains the general mechanism of body patterning, it does not provide detailed molecular mechanisms of as how the timing of a GRN is modulated by a morphogen gradient. Understanding of this conversion from temporal to spatial expression is essential for understanding the gene regulatory mechanisms driving patterning during development.

To answer this question, we focus in this study on gap genes in *Tribolium castaneum*, which offer a valuable system for understanding GRN dynamics during embryonic patterning. Gap genes were specifically selected because previous studies suggest their expression patterns closely follow the principles proposed by the speed regulation model (6,7,19,20,25–28). Specifically, gap genes *hunchback* (*hb*) (29,30), *Krüppel* (*Kr*) (31), *milles-pattes* (*mlpt*) (32), and *giant* (*gt*) (33) exhibit sequential waves of expression propagating from posterior to anterior regions, mirroring the temporal progression predicted by the speed regulation model (6). According to this model, a posterior-to-anterior gradient, namely Caudal (Cad) and/or Wnt signaling, controls the timing of sequential activation of gap genes, converting a temporal sequence of gene activation into a spatial pattern (6). Supporting this, RNA interference (RNAi) experiments that target parts of the Wnt pathway cause specific changes in both timing and positioning of gap genes expression. These changes fit what the speed regulation model predicts. However, it is not clear yet how Wnt/Cad modulates the overall timing of gap gene activation in Tribolium. Two hypotheses have been proposed to address this gap (6,19,34). The first (which we call the “general kinetic modulator” model) posits that the Wnt/Caudal gradient modulates specific kinetic factors that control the overall timing of the gap GRN, e.g. jointly modulating transcription and decay rates of multiple gap genes to control the speed of their activities based on Wnt/Cad concentrations. The second, encapsulated by what we term “GRN switching” model, suggests that the gradient does not directly influence transcription rates or decay rates, but rather modulates the wiring of GRN. This model proposes that gap genes are wired into two different GRNs. The first is a dynamic GRN, which is a genetic cascade that mediates the sequential activation of gap genes, where early-expressed genes, such as *hb*, activate subsequent targets like *Kr*, which in turn activates later genes such as *mlpt* and mediates the repression of earlier gene *hb*, and so on. The dynamic GRN is activated by high levels of the speed regulator Wnt/Cad, expressed in a posterior-to-anterior gradient in the Tribolium embryo. At lower levels of Wnt/Cad, gap genes are rather wired into a different network configuration: a static GRN, which is basically a multi-stable network that mediates the stabilization of gap gene expressions into stable expression domains.

To differentiate between the ‘general kinetic modulation’ model and the ‘GRN switching’ model, we employed HCR in situ hybridization (35), parental RNAi, and computational modeling to examine the dynamics and regulatory interactions of gap genes during AP patterning in both wild-type embryo and embryos in which gap genes are perturbed. Computational modeling suggests that the general kinetic modulation model predicts posterior-to-anterior propagation of gap gene expressions to be independent of genetic interactions. In contrast, the GRN switching model posits that this propagation depends on a transition in regulatory architecture, from a dynamic to a static gene regulatory network, and hence, dependent on GRN wirings.

So far, the genetic cascade hypothesis for gap genes has largely rested on two lines of evidence: the sequential activation of gap genes during early development (6,19), and phenotypic outcomes of gene knockdowns that (36), when synthesized across multiple studies, appear consistent with cascade-like regulation (6). However, these knockdown studies relied on late-stage expression patterns, after stabilization of gap gene domains, making it unclear whether the reported phenotypes reflected disruptions of the initial activation phase in the posterior growth zone or the stabilization phase in the anterior. Our results now resolve this issue. By tracking gap gene expressions in space and time across wild-type and perturbed embryos, we show that gap genes are indeed wired into a genetic cascade during the initialization phase in the posterior. This dynamic cascade drives sequential activation and propagation of expression waves.

Our experimental data further reveal that *shaven-baby* (*svb*), which functions together with *mlpt* as a gap gene in *Tribolium* (37), is initiated in the posterior as part of the dynamic cascade but later stabilized more anteriorly by the static GRN. In wild-type embryos, *svb* propagates anteriorly, while in *mlpt* RNAi embryos, svb fails to propagate and remains restricted to the posterior growth zone. This observation, consistent with the GRN switching model, indicates that propagation depends on dynamic cascade wiring, while anterior stabilization of svb relies on static regulatory interactions.

By presenting these findings, our study clarifies the phase-specific roles of gap gene GRNs and demonstrates that morphogen-controlled switching between dynamic and static network architectures is essential for robust spatiotemporal patterning.

## Results

### Spatiotemporal dynamics of gap gene expressions in WT Tribolium embryos

The spatiotemporal dynamics of gap gene expressions in *Tribolium* embryos have previously been described (6,30–33); however, we document them here again using HCR in situ hybridization to directly compare wild-type expression patterns with those observed following RNAi knockdown of specific gap genes (Fig. 1). In general, gap genes in *Tribolium* are expressed in a sequential manner, forming dynamic waves that originate in the posterior and propagate toward the anterior during development. To capture these dynamics, we analyzed gene expression at three-hour intervals from 14-17 hours AEL through 35-38 hours AEL (Fig. 1).

**Fig. 1.**
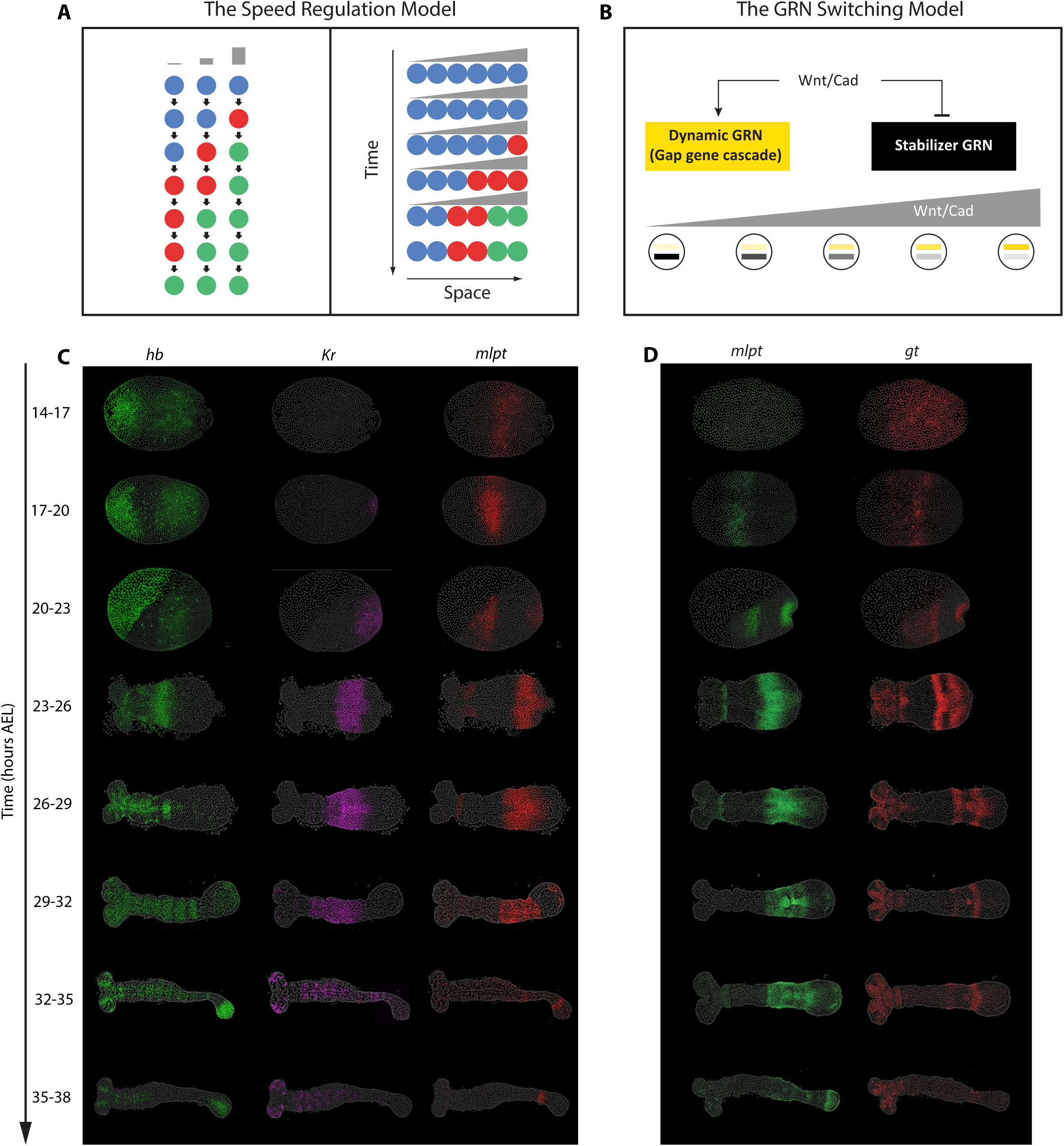
Models of gap gene regulation and spatiotemporal dynamics of gap gene expression in WT *Tribolium* embryos. (A) The Speed Regulation model. In this framework, a morphogen gradient modulates the speed of a temporal sequence of gene activations, such that sequential waves of expression are progressively frozen into spatial domains. (B) The GRN Switching model. Here, high Wnt/Cad levels activate a dynamic gene regulatory network (GRN) in the posterior, organized as a genetic cascade, while lower Wnt/Cad levels anteriorly favor a static stabilizer GRN that locks gene expression into stable domains. (C, D) FISH detecting subsets of the four core gap genes (*hb, Kr, mlpt, gt*) in staged wild-type embryos (14-38 h AEL). Panel C shows expression of *hb*, *Kr*, and *mlpt*, while Panel D shows *mlpt* and *gt*. Gene expressions are visualized in the corresponding channels/colors as indicated in the figure. All embryos are oriented with anterior to the left and posterior to the right.

During the blastoderm stages (14-17, 17-20, 20-23 h AEL), zygotic *hb* initiates in the serosa and posterior, then clears from the posterior to leave a stable anterior domain; during the germband phase (23-26 to 32-35 h AEL) the posterior remains devoid of *hb* until 32-35 h, when posterior expression reappears and persists through AP patterning (Fig. 1A, green). *Kr* emerges at 17-20 h AEL at the posterior of the blastoderm, clears from posterior by 23-26 h AEL leaving an expression domain more anteriorly, and remains absent from the posterior thereafter (Fig. 1A, purple). *Mlpt* expression begins at 20-23 h AEL in the posterior, clears between 23-26 and 26-29 h leaving an anterior domain, and at 29-32 h is re-expressed posteriorly but fades rapidly to a narrow posterior stripe. (Fig. 1A, red and 1B green). *gt* is first detectable at 20-23 h AEL, undergoes rapid clearing and reappearing, resulting in two distinct stripes by 26-29 h AEL (Fig. 1B, red). All resolved gap gene expression patterns at the anterior persist for a while, before eventually fading, except for the most posterior late expressions of *hb*, which stay restricted to the growth-zone. Some genes are also activated in additional domains that are not considered in this study. For example, *hb* and *Kr* are expressed in the CNS of anterior germbands (*hb* from 29–32 h AEL onward, and *Kr* from 32– 35 h AEL onward). In addition, *hb* is expressed in the serosa, while *mlpt* and *gt* are activated in the head primordia (Fig. 1).

Taken together, these gene expression patterns illustrate a general principle of *Tribolium* gap-gene regulation: gap genes are activated sequentially in dynamic waves that initiate in the posterior and move anteriorly. This posterior-to-anterior propagation underlies the temporal unfolding of the body-plan during early embryogenesis.

### Knockdown phenotypes of gap genes suggest that they are wired into a genetic cascade

The sequential activation of gap genes raises the possibility that these genes may be organized into a genetic cascade. Previous studies describing various *Tribolium* gap gene knockdown phenotypes (30–33,36) have previously led us to the conclusion that gap genes might indeed function as a genetic cascade (6). However, those previous observations provided only static snapshots at specific developmental stages, lacking detailed temporal resolution. Without dynamic information, phenotypes observed in knockdown embryos might only superficially resemble those expected from a genetic cascade, rather than truly reflecting sequential genetic interactions. Because a genetic cascade inherently unfolds over time, temporal information is essential to rigorously test the genetic cascade hypothesis.

Therefore, we systematically tracked the spatiotemporal dynamics of gap gene expressions following individual gap-gene knockdowns, using carefully staged egg collections. Below, we describe how the sequential activation of gap genes and the associated propagating waves are affected by specific gap gene disruptions.

In a genetic cascade, each gene’s expression is typically expected to mediate activation of the subsequent gene, while simultaneously facilitating the repression of preceding genes. Notably, this cascade arrangement does not necessarily imply direct positive activations. A cascade can also be composed entirely of inhibitory interactions, with sequential activation arising indirectly through de-repression (i.e., repression of repressors). Regardless of the exact mechanism, both genetic cascade configurations predict that knocking down one gene should disrupt the expression of genes that are normally activated later in the cascade, while causing persistent or expanded expression of genes that act earlier in the sequence. Here, we directly test these predictions by examining gap gene expression dynamics across both space and time following targeted RNAi knockdowns.

#### *hb* RNAi disrupts downstream gap-gene expression

To examine the role of *hb* within the gap GRN, we performed parental RNAi targeting *hb* and analyzed embryos from 14-17 to 32-35 h AEL (Fig. 2). In *hb* RNAi embryos, *hb* expression was absent or strongly reduced at all stages examined, confirming efficient knockdown (Fig. 2A).

**Fig. 2.**
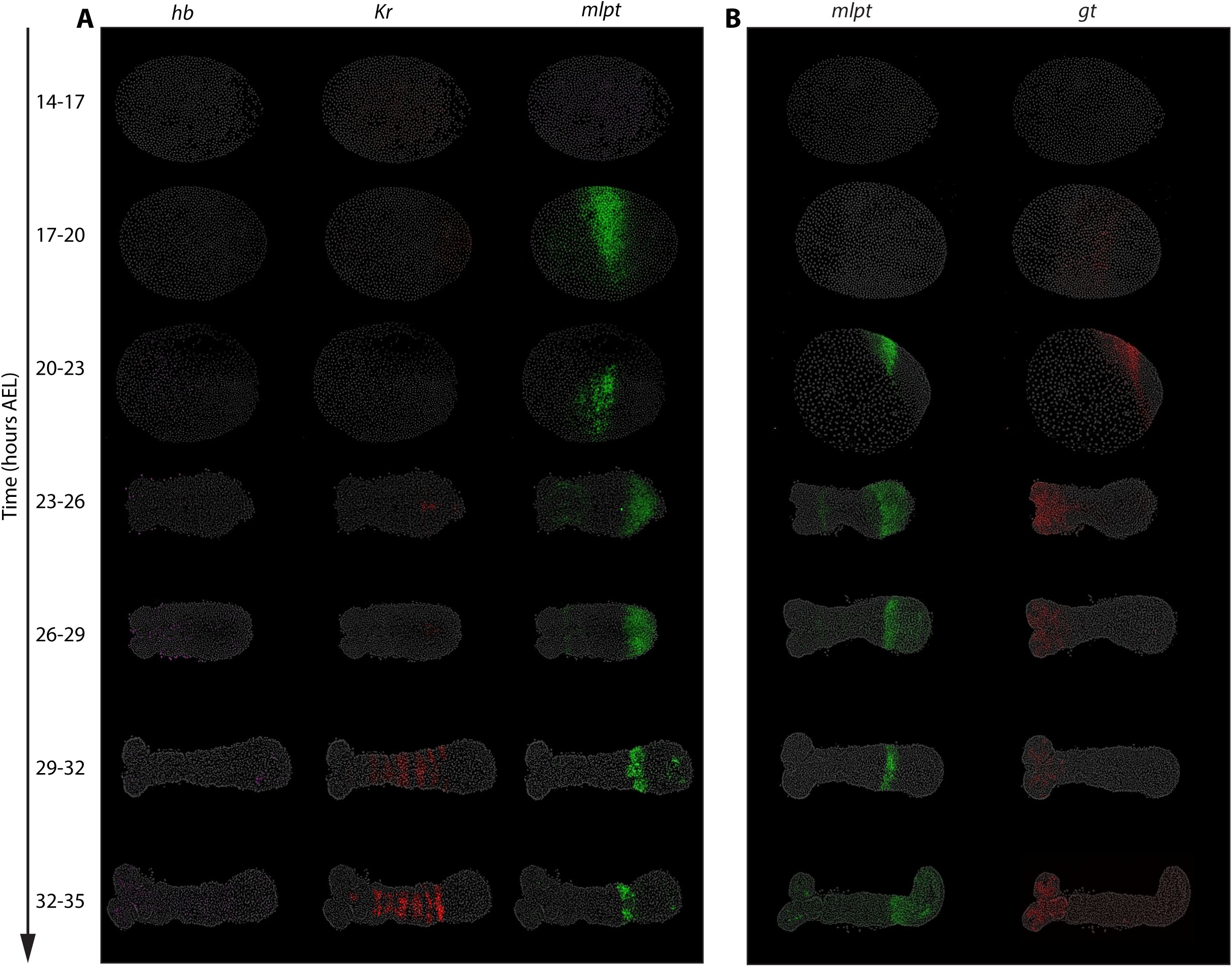
*hb* RNAi disrupts downstream gap-gene expression. Panels A and B show fluorescence in situ hybridization (FISH) detecting subsets of the core gap genes (*hunchback, Krüppel, milles-pattes, giant*) in staged embryos (14-35h AEL). Panel A displays the expression patterns of *hb, Kr, and mlpt*, while Panel B shows the expression of *mlpt* and *gt.* Gene expressions are visualized in the corresponding channels/colors as indicated in the figure. All embryos are oriented with anterior to the left and posterior to the right.

Consistent with *hb* acting at the top of the gap cascade, the trunk expressions of subsequent gap genes in the cascade were mostly diminished: *Kr* (Fig. 2A, red; except for the late striped *Kr* expression in the CNS), most of *mlpt* expression (Figs. 2A; 2B, green), and *gt* (Fig. 2B, red). Notable exception is the expressions of *mlpt*. A small anterior subdomain of the early posterior *mlpt* domain persisted in *hb* RNAi embryos, and thus appears to activate independently of other gap inputs (Figs. 2A; 2B, green).

Notably, Kr, (most of) *mlpt*, and *gt* expressions in *hb* knockdowns fail to initiate in the posterior, rather than appearing and later being lost after anterior stabilization. This supports the view that gap genes are wired into a genetic cascade in the posterior, where high Cad levels drive their sequential activation, and highlights *hb* as the initiating gene required to trigger this cascade during early embryonic patterning.

#### RNAi knockdown of *Kr* results in the expansion of *hb* expression domain and the reduction of *mlpt* and *gt* expressions

In embryos subjected to *Kr* RNAi, *Kr* expression was indistinguishable from background levels in most embryos, confirming effective knockdown (Fig. 3A; 3B, purple). Under these conditions, the *hb* expression domain expanded anteriorly, consistent with *Kr* normally repressing its upstream gene *hb* (Fig. 3A, green). Conversely, genes that normally act later in the cascade showed severely reduced expression. The *mlpt* domain was either strongly diminished or completely absent, except for a small anterior region of expression, which again appears to be activated independently of earlier genes in the cascade (Fig. 3A, red and 3B, green).

**Fig. 3.**
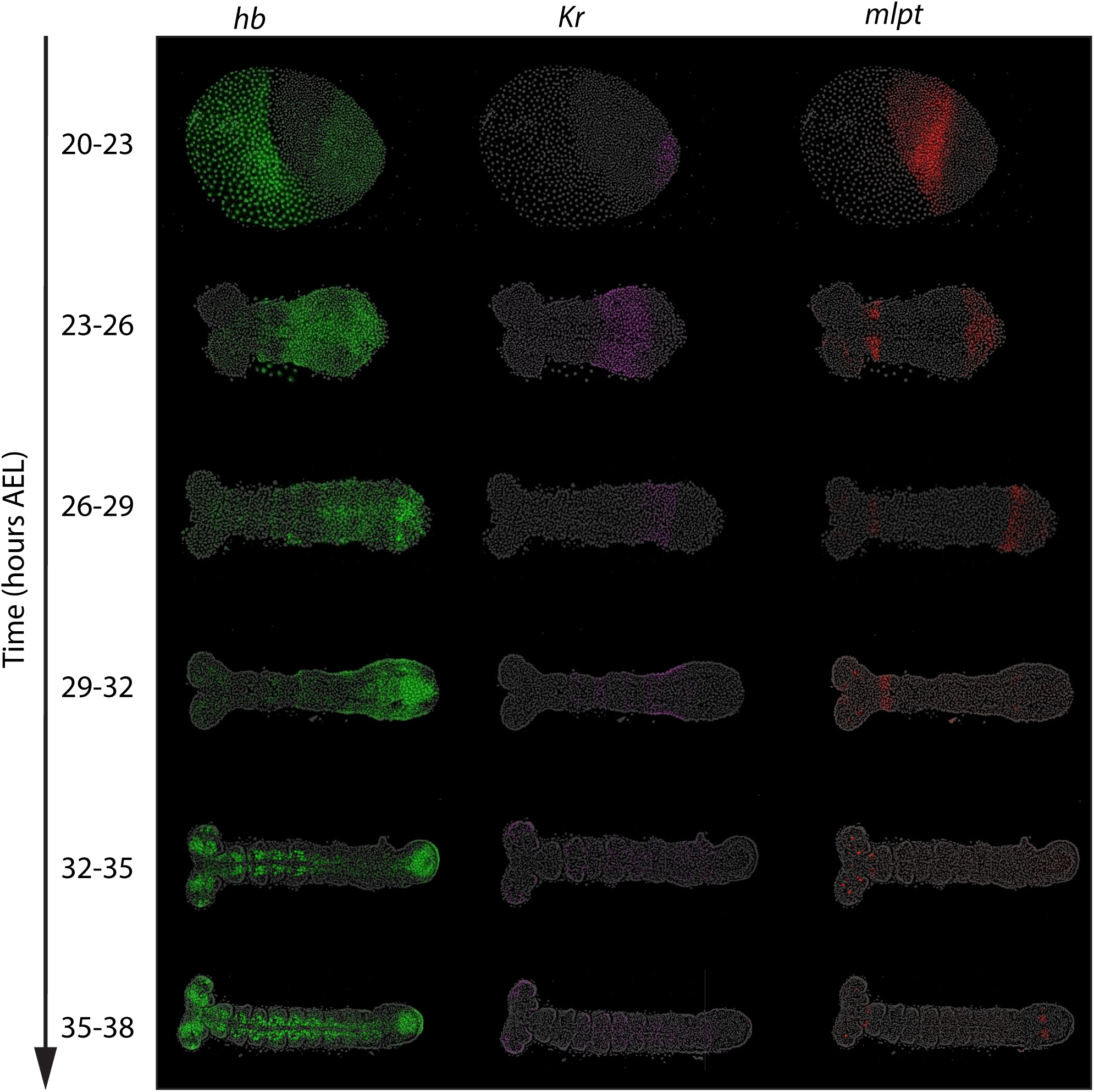
*Kr* RNAi expands *hb* expression domain and reduces *mlpt* expressions. Panels A and B show fluorescence in situ hybridization (FISH) detecting subsets of the core gap genes (*hunchback, Krüppel, milles-pattes*) in staged embryos (20–38 h AEL) following parental *Kr* RNAi. Gene expressions are visualized in the corresponding channels/colors as indicated in the figure. All embryos are oriented with anterior to the left and posterior to the right

Notably, genes that act later in the cascade than *Kr* failed to initiate altogether, rather than first appearing posteriorly and then disappearing after their stabilization in anterior domains. At the same time, the earlier gene *hb* remained persistently expressed in the posterior upon *Kr* knockdown, instead of being cleared from the posterior initially and then expanding toward the posterior after stabilization. These results confirm the regulatory role of *Kr* within the gap GRN and align with a gap genetic cascade model active in the posterior, where each gene both promotes the activation of the next in sequence and mediates the repression of the one expressed earlier.

#### *mlpt* RNAi expands *Kr* posteriorly and abolishes *gt* expression

Embryos subjected to parental RNAi targeting *mlpt* showed a complete loss of *mlpt* expression, indicating successful knockdown (Figs. 4A, red; 4B, green). Under this condition, *Kr* expression fails to clear from the posterior at 23-26 h AEL and 26-29 h AEL, where it does clear in the WT (compare with Fig. 1), resulting in the extension of *Kr* expression significantly toward the posterior where it remained broader compared to wild-type embryos, suggesting that *mlpt* normally represses *Kr* expression (Fig. 4A, magenta). In contrast, the trunk *gt* expression was completely absent in *mlpt* RNAi embryos (Fig. 4B), implying that *mlpt* is necessary for the activation of *gt*.

**Figure 4.**
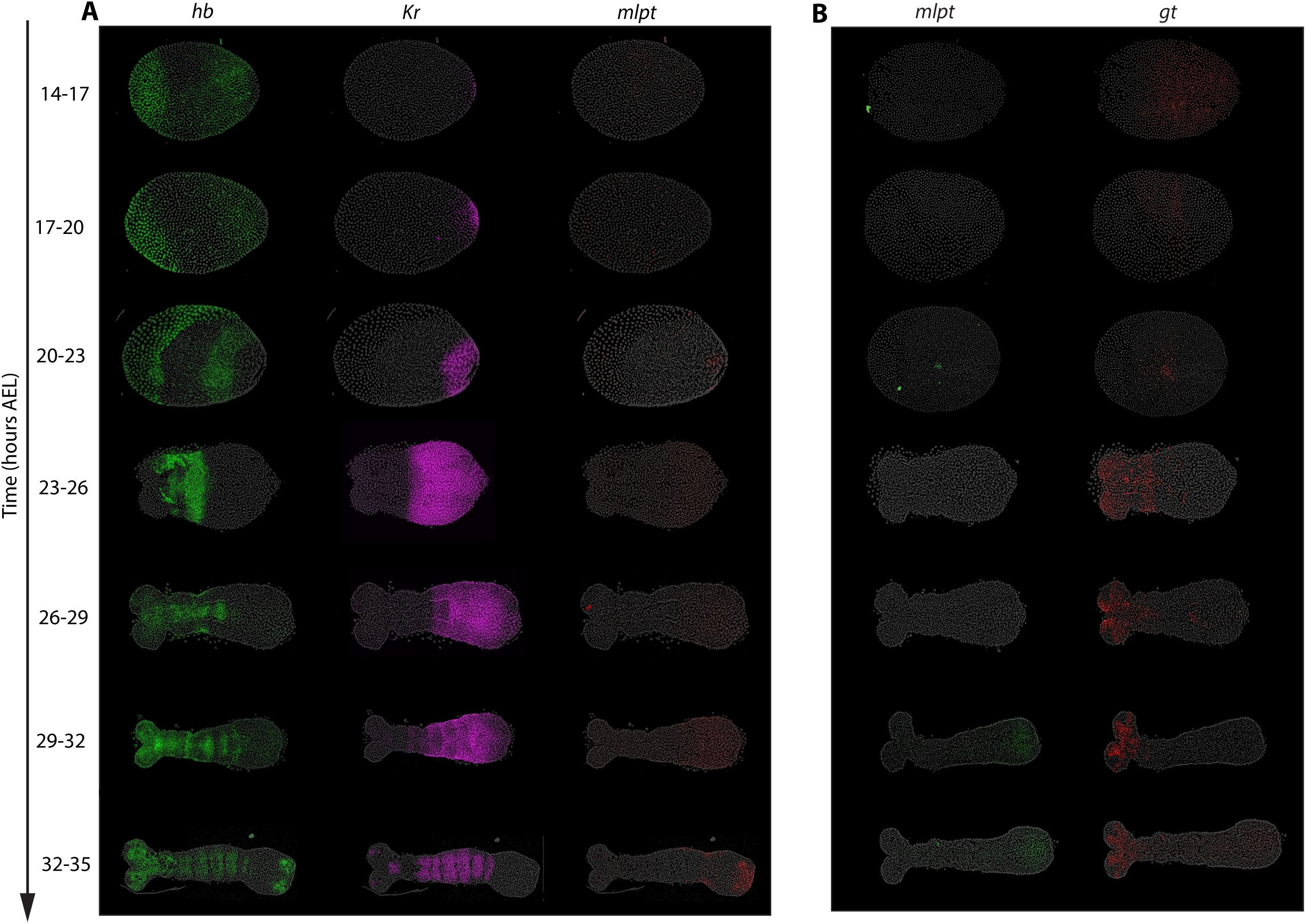
*mlpt* RNAi expands *Kr* posteriorly and abolishes *gt* expression. Panels A and B show fluorescence in situ hybridization (FISH) detecting subsets of the core gap genes (*hunchback, Krüppel, milles-pattes, giant*) following parental *mlpt* RNAi in staged embryos (14-35h AEL). Panel A displays the expression patterns of *hb*, *Kr* and *mlpt*. Panel B shows the expression of *mlpt* and *gt*. Gene expression is visualized in the corresponding channels/colors as indicated in the figures. All embryos are oriented with anterior to the left and posterior to the right.

Notably, in *mlpt* RNAi embryos, *gt* failed to initiate altogether, rather than appearing posteriorly and then failing to stabilize, while *Kr* remained persistently expressed in the posterior instead of being cleared and later expanding more anteriorly during the stabilization phase. These findings confirm the regulatory role of *mlpt* within the gap gene network and support its position as an integral component of a genetic cascade that operates in the posterior during the initialization phase of gap gene expression, where it mediates both the activation of the next gene (*gt*) and the repression the preceding gene (*Kr*).

#### *gt* RNAi results in posterior expansion of *mlpt* expression

In embryos subjected to *gt* RNAi, *gt* expression was reduced to background levels from 14–17 h AEL onward, confirming effective knockdown (Fig. 5A, red). Under these conditions, *mlpt* expression failed to clear from the posterior at either 23–26 h AEL or 26–29 h AEL, when it normally does in wild type (Fig. 1). As a result, *mlpt* persisted in the posterior and its domain expanded beyond normal boundaries (Fig. 5B, red; Fig. 5A, green), indicating that *gt* normally represses *mlpt*. Together, these observations identify *gt* as a negative regulator of *mlpt* within a genetic cascade that operates in the posterior during the initialization phase of gap gene expression.

**Figure 5.**
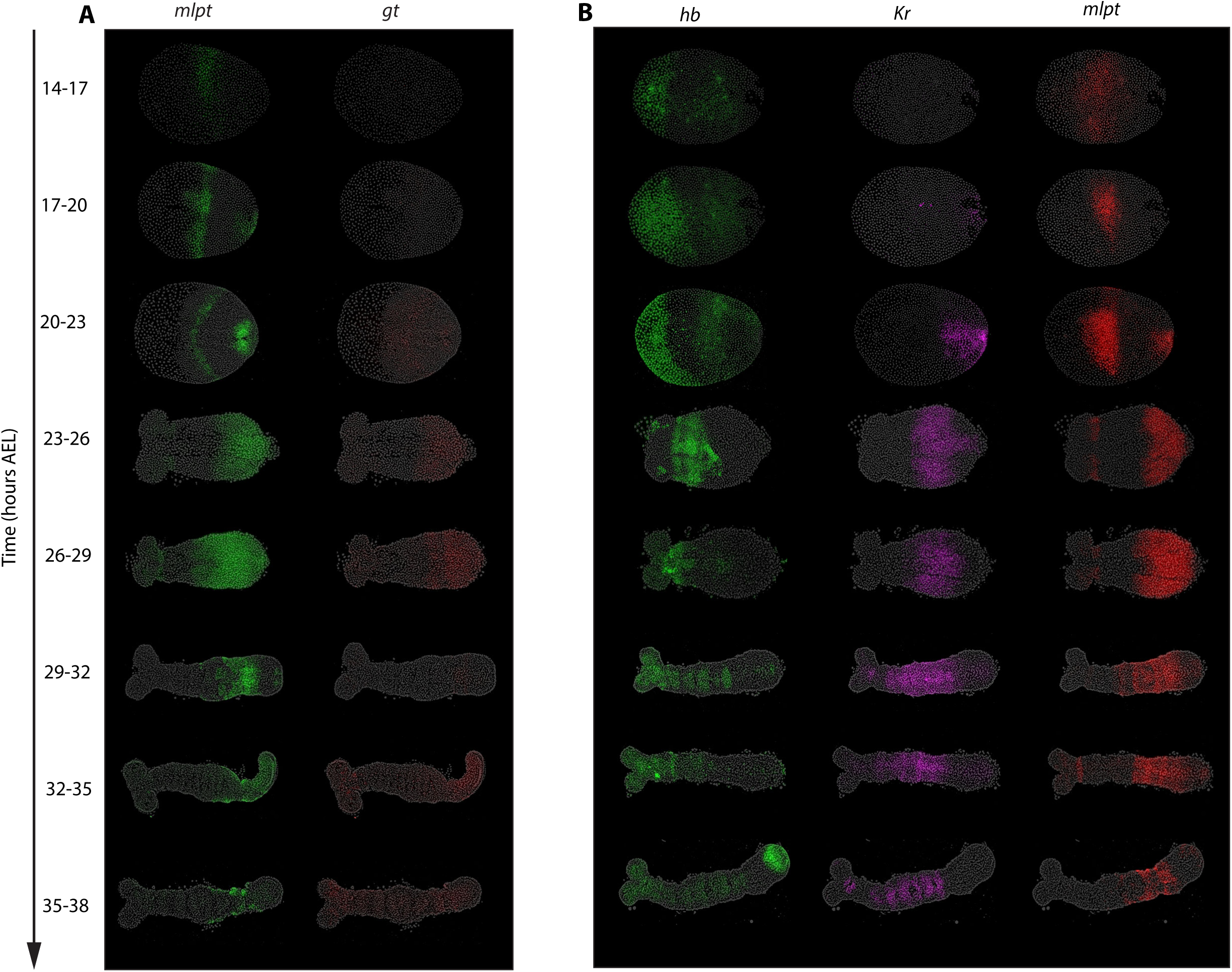
*gt* RNAi results in posterior expansion of *mlpt* expression. Panels A and B show fluorescence in situ hybridization (FISH) detecting subsets of the core gap genes (*hunchback, Krüppel, milles-pattes, giant*) following parental *gt* RNAi in staged embryos (14-38h AEL). Panel A shows the expression patterns of *mlpt* and *gt*. Panel B displays the expression of *hb*, *Kr*, and *mlpt*. Gene expressions are visualized in the corresponding channels/colors as indicated in the figure. Embryos are oriented with posterior to the right in all panels.

### The role and regulation of svb within the gap genetic cascade

*shavenbaby (svb)* encodes a transcription factor best known in *Drosophila* for controlling epidermal differentiation, where it regulates the formation of cuticular trichomes (38,39). In its default state, Svb acts as a transcriptional repressor. Its activity is modulated by small peptides encoded by the *mlpt* locus, which bind Svb and induce a conformational change that converts it into a transcriptional activator. This Svb-Mlpt module is an evolutionarily conserved switch controlling diverse processes, from epidermal differentiation to embryonic segmentation. In *Tribolium*, both genes are expressed dynamically during early development, placing them in a position to function as part of the gap gene network (37).

#### Mlpt-bound Svb activates *gt* expression

In *Tribolium*, *svb* first appears as a posterior cap at 20-23 h AEL, then clears from the posterior, leaving behind a stripe of expression more anteriorly. It is subsequently re-expressed and maintained at the posterior through later patterning stages (Fig. 6C: *svb* in WT). Our data reveal a critical regulatory interaction between *svb* and *mlpt* during embryogenesis that is essential for proper gap gene expression. Consistent with their known biochemical interaction, we find that the Mlpt-bound Svb complex activates target genes, notably *giant (gt)*. RNAi experiments confirm this: embryos lacking either *mlpt* or *svb* fail to express *gt* (Fig. 4B, red; Fig. 6B, red). These observations place *mlpt* and *svb* as cooperative components of the posterior genetic cascade, linking their conserved biochemical partnership to the regulation of early AP patterning genes.

**Figure 6.**
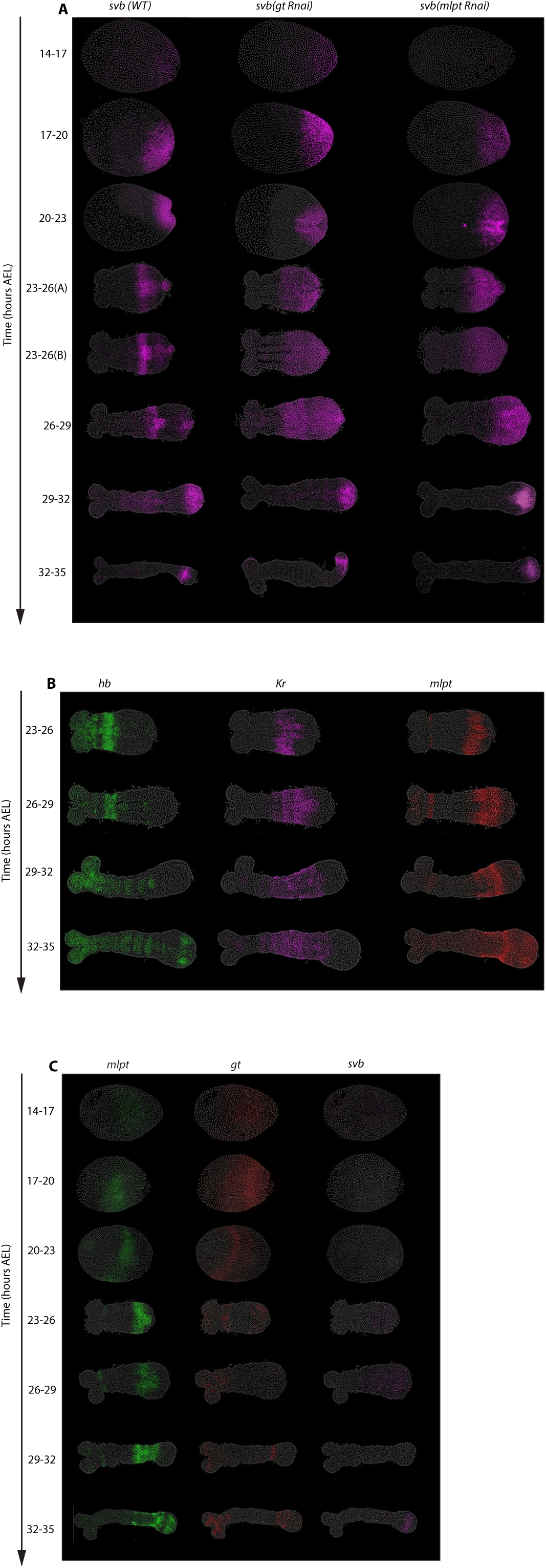
The role and regulation of *svb* within the gap genetic cascade. Panel A (23–32h AEL) shows the expression patterns of *hb, Kr,* and *mlpt* following parental *svb* RNAi. Panel B (14–35h AEL) displays the expression of mlpt, *svb and gt*. Panel C shows the expression of *svb* wild-type (WT), *mlpt RNAi*, and *gt RNAi* embryos. Embryos were analyzed at various developmental stages (14–IT35h AEL). All embryos are oriented with anterior to the left and posterior to the right.

#### *mlpt* mediates the transition from dynamic to static phases of *svb* expression

An unresolved question concerns the regulatory mechanisms controlling *svb* expression itself. As described above, *svb* expression initially emerges as a posterior cap at the blastoderm stage (20-23 h AEL), closely mirroring the expression domain of *cad* (6,12,40–42). Upon activation of *mlpt*, the posterior *svb* expression clears, and a distinct anterior expression band forms (Fig. 6C). Crucially, in *mlpt* RNAi embryos, *svb* expression remains trapped within the posterior growth zone, failing to propagate anteriorly (Fig. 6C). This highlights a critical role for *mlpt* in facilitating anterior propagation of *svb*. Importantly, *mlpt* cannot drive this transition on its own, since it does not encode a transcription factor; rather, it exerts its function specifically through interaction with *svb*, enabling Svb to act as an activator.

Thus, the most parsimonious model for stabilizing *svb* expression outside the growth zone involves the autoactivation of *svb* mediated by the *mlpt*-bound *svb* complex. For other genes, direct autoactivation is difficult to experimentally confirm, since knocking down the gene disrupts its expression entirely, obscuring any specific autoactivation link. However, in the case of *svb*, autoactivation is uniquely detectable through *mlpt* RNAi. Because *mlpt* is necessary for Svb’s conversion into a coactivator, targeting *mlpt* selectively abolishes the autoactivation feedback loop without eliminating all *svb* expression, revealing the presence of this regulatory mechanism.

#### *gt* mediates the clearance of *svb* from the growth zone

In wild-type embryos, posterior clearance of *svb* occurs around 23-26 h AEL, likely mediated through repression by *gt* (Fig. 6C). In *gt* RNAi embryos, *svb* successfully propagates anteriorly out of the growth zone, presumably because *mlpt* expression remains intact, but fails to clear from the posterior growth zone (Fig. 6C).

Collectively, these results show that the anterior propagation of *svb* expression depends on the Svb-Mlpt complex, which drives autoactivation of *svb* within a stabilizing GRN operating outside the growth zone. This is consistent with the GRN switching model, in which the stabilization phase is mediated by a static network that takes over from the dynamic cascade active in the posterior growth zone, thereby ensuring the transition from sequential activation to stable expression domains.

### Computational modeling supports GRN switching as the mechanism for spatial stabilization of gap gene expression patterns in *Tribolium*

To investigate whether our experimental observations are consistent with the GRN switching model, we developed a computational framework based on coupled ordinary differential equations (ODEs) that encode the gap gene regulatory interactions inferred from our experiments (Fig. 7). The model incorporates the six gap genes studied here (*hb, Kr, mlpt, svb, gt,* and a putative regulator X) and explicitly represents the dynamic module active in the posterior and the static module stabilizing expression anteriorly. Critically, it also includes the Mlpt–Svb interaction, which our experiments identify as essential for activating *gt* and maintaining autoactivation of *svb*.

**Figure 7.**
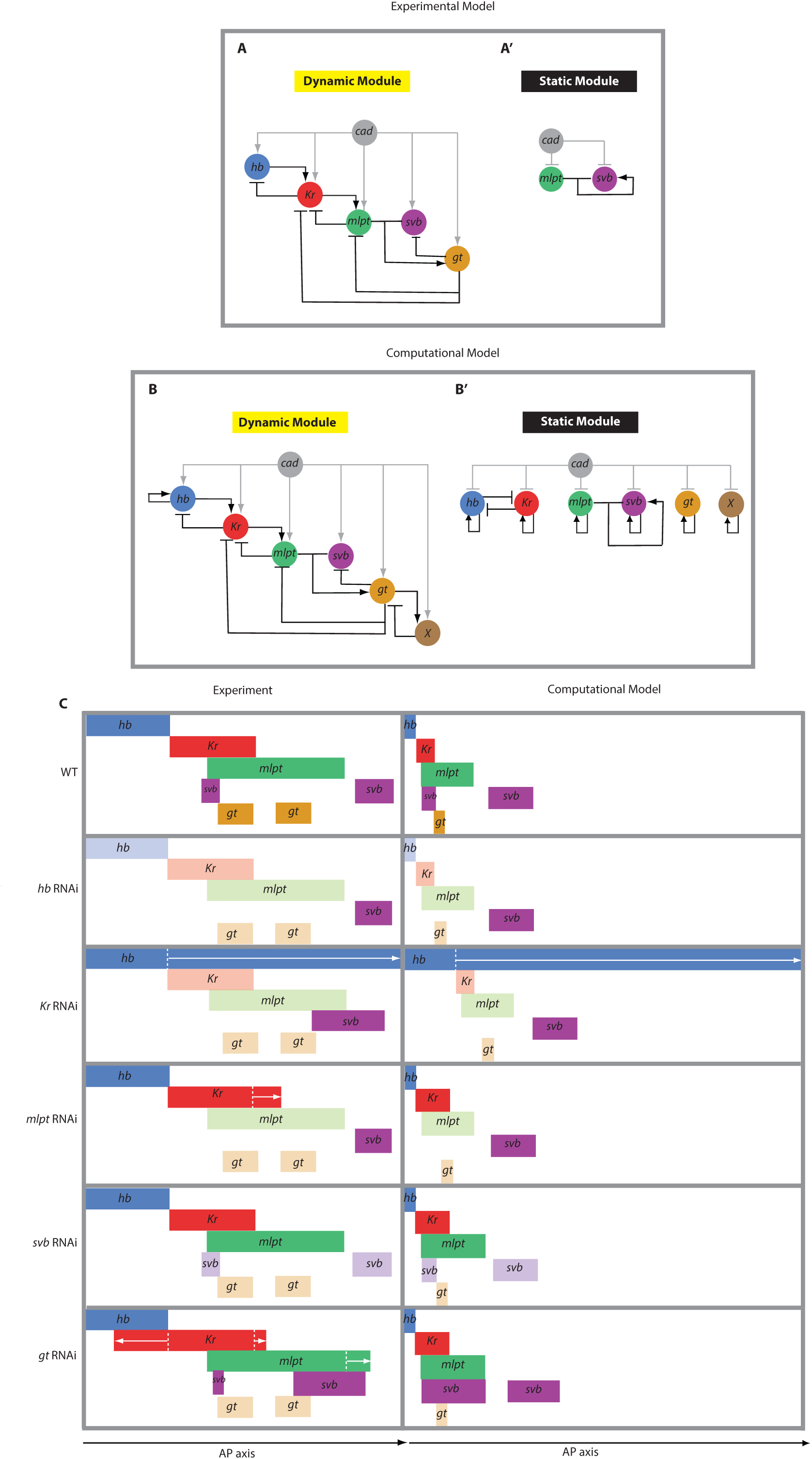

#### Computational model and *in silico* RNAi

The computational model (Fig. 7B) integrates gene expression dynamics governed by interactions informed by our RNAi data (Fig. 7A). Genes are modeled through coupled ODEs, including terms for transcriptional activation, repression, and first-order mRNA degradation. The dynamic module (Cad-dependent) mediates the sequential activation of gap genes in the posterior, whereas the static module maintains stabilized anterior gene expression domains independent of Cad. An in silico RNAi option was included by explicitly setting expression and transcriptional derivatives of targeted genes to zero, enabling precise comparison to experimentally observed RNAi phenotypes.

#### Wild-type simulation matches observed spatiotemporal dynamics of *svb* and *gt*

In wild-type simulations (Fig. 7C), *svb* initially mirrors the posterior-to-anterior gradient of *Cad*, similar to observed embryonic data. Subsequently, upon expression of *mlpt*, formation of the *mlpt-svb* complex establishes a positive auto-regulatory loop that propagates *svb* anteriorly, beyond the original posterior growth zone. Concurrently, posterior expression of *gt*, activated by the *mlpt-svb* complex, represses *svb* within the growth zone, resulting in posterior clearance of *svb* expression. Thus, the model captures key experimental observations: initial posterior *svb* activation by *Cad*, anterior propagation mediated by mlpt via auto-activation, and posterior clearance through repression by *gt*. Additionally, the model accurately replicates the experimentally observed repression of *mlpt* and *svb* within the posterior growth zone by *gt*, underscoring its role as a negative regulator essential for maintaining sharp boundaries of gene expression domains.

#### in silico *hb* and *Kr* RNAi phenotypes mirror those observed experimentally

The computational model successfully recapitulates the experimental phenotypes observed in *hb* and *Kr* RNAi embryos. When *hb* was knocked down in silico, expression of later-acting genes in the cascade (*Kr, mlpt,* and *gt*) failed to initiate in the posterior, while a small anterior domain of *mlpt* persisted independently of *hb*. This outcome matches our experimental findings (Fig. 7C) and confirms the role of *hb* as the initiating gene in the cascade.

Similarly, in silico knockdown of *Kr* reproduced the dual effect observed experimentally (Fig. 7C). The expression of *hb* expanded anteriorly, consistent with *Kr* normally repressing *hb*. At the same time, *mlpt* and *gt* expression failed to arise, except for a small anterior domain of *mlpt* that appears to be activated independently of the cascade. Together, these simulations closely mirror the observed RNAi phenotypes and further validate the cascade wiring inferred from experimental perturbations.

#### in silico *mlpt* RNAi restricts *svb* expression to the growth zone, mirroring experimental observations

Simulated RNAi targeting *mlpt* reveals critical insights consistent with our RNAi experiments (Fig. 7C). With *mlpt* knocked down in silico, the absence of the *mlpt-svb* complex abolishes the positive feedback loop necessary for anterior propagation of *svb* expression. Consequently, in silico *svb* expression remains confined within the Cad-rich posterior growth-zone throughout development, precisely mirroring the experimentally observed phenotype in *mlpt* RNAi embryos (Fig. 7C). The modeling thus confirms our interpretation that anterior propagation of *svb* relies specifically on the *mlpt-svb* interaction, reinforcing the proposed regulatory role of *mlpt* within the GRN.

#### in silico *gt* RNAi broadens the *mlpt* expression domain and disrupts posterior clearance of *svb*

Simulated *gt* RNAi further supports our experimental findings, revealing two key outcomes. First, the absence of *gt*-mediated repression results in prolonged persistence within the growth zone and the expansion of the *mlpt* expression domain towards posterior. Second, without posterior repression by *gt*, *svb* fails to clear from the growth zone but continues to propagate and stabilize anteriorly through the intact *mlpt-svb* auto-regulatory feedback loop. Thus, the computational *gt* RNAi scenario closely matches experimentally observed phenotypes, where the removal of *gt* leads to persistent expression of *mlpt* and *svb* in the posterior.

A noteworthy comparison arises between *svb* expression in *mlpt* versus *gt* RNAi conditions, observed both experimentally and in silico (Figs. 6 and 7). In both cases, *svb* fails to clear from the posterior growth zone. However, the anterior dynamics differ: in *mlpt* RNAi embryos, *svb* is unable to propagate anteriorly, whereas in *gt* RNAi embryos, *svb* successfully propagates anteriorly but remains abnormally persistent in the posterior, producing an expanded expression domain. This distinction can be explained by the requirement of *mlpt* for anterior propagation of *svb*: in *gt* RNAi embryos, *mlpt* is still expressed, enabling propagation, while in *mlpt* RNAi embryos, its absence prevents anterior spread.

#### Implications: Computational results strongly support GRN switching

Our computational modeling shows that the experimental phenotypes in *Tribolium* are best explained by the GRN switching model. While the “general kinetic modulator” model can, in principle, generate stable gene expression domains, it does so without requiring genetic interactions for stabilization: domain stability arises automatically from the decline of the kinetic modulator. This prediction does not match our experimental findings. In particular, *svb* fails to propagate anteriorly in *mlpt* RNAi embryos, showing that stabilization depends on the Mlpt-Svb interaction, a feature the general kinetic modulator model cannot account for. In contrast, the GRN switching framework explicitly incorporates a morphogen-dependent transition from a dynamic cascade to a static stabilizing network, and thus correctly predicts this behavior. Importantly, our computational simulations reproduce all observed phenotypes in wild type and gap gene knockdowns, including the contrast between *svb* expression in wild type, *mlpt* RNAi, and *gt* RNAi. Together, these results indicate that GRN switching provides a more accurate explanation of gap gene dynamics in *Tribolium* than the general kinetic modulator model.

## Discussion

Embryonic patterning is the process by which cells acquire distinct fates in precise spatial arrangements, enabling the formation of complex body plans. Classical frameworks, such as the French Flag model, have emphasized spatial control, where morphogen gradients are interpreted by cells as positional information to establish domains of gene expression. More recently, however, evidence has accumulated that temporal dynamics also play a central role in pattern formation (7–9,17,25,43–50). In several developmental contexts, transcription factors are activated in a defined sequence over time, a logic termed temporal patterning (51,52). In neuroblast lineages, such temporal cascades diversify neuronal fates without directly mapping onto spatial order, because progenitors delaminate and migrate (52–55). By contrast, in other tissues (including vertebrate Hox regulation, somitogenesis, and insect segmentation) temporal cascades appear to be translated directly into spatial registers, such that the order of gene activation in time is preserved in space (7). This emerging concept, that embryonic patterning can involve a conversion of temporal order into spatial pattern, raises a central question: what molecular mechanisms allow dynamic gene regulatory programs to be “frozen” into stable spatial domains?

Here we have addressed this question using the gap gene network in *Tribolium castaneum* as a case study. Gap genes have been suggested to act in a genetic cascade, based on their sequential activation and on RNAi phenotypes that appeared consistent with cascade-like regulation (6,20). However, these inferences were drawn from static, late-stage expression patterns, when anterior domains were already stabilized, making it unclear whether a true cascade operates during the initial posterior activation phase (6,36). By directly following gap gene dynamics, we find that gap genes indeed participate in a genetic cascade, but only during the posterior initialization phase. Our data are consistent with a model in which the gap gene system operates in two distinct phases: a dynamic genetic cascade driving posterior activation, followed by a static GRN that mediates the propagation of gap gene expressions towards anterior and their stabilization into spatial expression domains.

This dual organization offers a mechanistic explanation for how temporal sequences are translated into spatial patterns. In the posterior, a genetic cascade composed of gap genes mediates their sequential activation, while in the anterior, a static GRN stabilizes these expressions into fixed domains. The regulation of *svb* provides a clear example (Fig. 6): its expression begins in the posterior and propagates anteriorly upon *mlpt* activation in wild type embryos. In *mlpt* knockdowns, this propagation fails, showing that stabilization of anterior expression depends on specific genetic interactions rather than on global kinetic modulation. Computational modeling recapitulates these dynamics (Fig. 7), reinforcing the view that morphogen-dependent rewiring underlies the observed behaviors. The distinction between dynamic and static phases also clarifies why general kinetic modulation is insufficient. A model in which morphogen gradients globally alter transcription or decay rates can in principle change the pace of expression, but it cannot explain why the same set of genes function first as a cascade and then as a stabilizing network. GRN switching provides a stronger account: morphogen input alters not just kinetic parameters but the wiring of the network, thereby producing phase-specific behaviors.

This raises another important question: if the GRN switching scheme is correct, how can the same genes participate in two different networks: one dynamic at the posterior, the other static at tge anterior? One plausible explanation is that each gene is controlled by distinct enhancers that encode its wiring within each GRN, a mechanism formalized in the “Enhancer Switching” model (6,34). In this framework, every patterning gene possesses a “dynamic” enhancer, which mediates its role in the posterior cascade, and a “static” enhancer, which governs its stabilization in the anterior. Morphogen levels then determine the balance between these enhancers, promoting dynamic activity in the posterior and static activity in the anterior. The model successfully generates gene expression waves in silico (6), and is in line with observations in *Drosophila* (56,57) and vertebrate systems (58,59). While promising, the model necessitates rigorous testing to determine if refinement or alteration is needed. An alternative possibility is enhancer pleiotropy (60–62), in which a single enhancer encodes multiple regulatory logics, enabling a gene to participate in both phases through context-dependent outputs.

Recent technical advances make it possible to directly test these ideas in *Tribolium*. A framework combining ATAC-seq with MS2-based live enhancer reporters has been developed to discover active enhancers and follow their dynamics in vivo (63). Initial results from this system have revealed candidate enhancers with dynamic- or static-like activities, providing tentative support for the Enhancer Switching model, but stronger evidence will require systematic identification of enhancer pairs and functional tests through enhancer-specific deletions. Applying this framework to gap genes will be essential to determine whether their dual roles in posterior cascades and anterior stabilization are mediated by distinct enhancers or by pleiotropic regulatory logic.

Taken together, our findings establish the gap gene network as a mechanistic example of how temporal cascades can be translated into spatial domains. They suggest that GRN switching may represent a general principle for time-to-space conversion in development, and perhaps an evolutionary innovation that enabled temporal programs to be co-opted for spatial patterning during the transition to multicellularity (51). At the same time, they point forward to the enhancer level, where rigorous tests of enhancer function will be essential to determine how morphogens modulate the timing of GRNs at the molecular scale. By linking broad conceptual models to concrete molecular mechanisms, our study advances the emerging view that embryonic patterning is as much a problem of temporal regulation as of spatial information.

## Material and method

### Beetle cultures

Beetle cultures were reared on flour supplemented with 5% dried yeast in a temperature- and humidity-controlled room at 24 °C. To speed up development, beetles were reared at 32 °C.

### Egg collections for developmental time windows

Developmental time windows of 3 hr were generated by incubating 3 hr egg collections at 24 °C for the desired length of time. Beetles were reared in flour supplemented with 5% dried yeast.

### Fixation of Embryos

Tribolium castaneum eggs were collected on flour plates, floated off in 50 % bleach (∼2 min), rinsed in water, and fixed at room temperature for 1 h in a two-phase formaldehyde/heptane mixture. Devitellinization was achieved by rapid shaking with methanol, followed by repeated passage through a 20-gauge needle. Embryos were washed three times in methanol and stored at −20 °C.

### In situ hybridization of fixed embryos

Samples were rehydrated through 50 % MeOH/PBT into PBT (1 × PBS, 0.1 % Tween-20), post-fixed in 4 % formaldehyde/PBT for 30 min and rinsed in PBT. Embryos were pre-hybridized for 30 min in 30 % probe hybridization buffer at 37 °C, then incubated overnight with gene-specific HCR probes (Molecular Instruments) at 37 °C. After four 15-min washes in 30 % probe wash buffer (37 °C) and three 5-min washes in 5 × SSCT, snap-cooled fluorescent hairpins were applied in amplification buffer and allowed to react overnight at room temperature in the dark. Excess hairpins were removed by sequential washes in 5 × SSCT, and embryos were mounted in SlowFade™ Gold (Thermo Fisher).

### Imaging and image processing

Confocal Z-stacks were acquired on Leica TCS SP8 and Nikon AX systems. Maximum-intensity projections were generated in Fiji/ImageJ.

### dsRNA Injection

Parental RNAi was performed to knockdown various gap genes. For dsRNA synthesis, a plasmid containing portions of a gap gene CDS flanked by T7 and T3 sites was used as template. T7 and T7-T3 primers were used to amplify the gap genes CDS fragment using the CloneAmp HiFi PCR Premix (Takara Bio). PCR cycle conditions were as follows: denaturation at 98 °C for 15 sec; annealing at 40 °C (first 5 cycles)/ 55 °C (remaining 25 cycles) for 15 sec; elongation at 72 °C for 1 min. The obtained PCR fragment was purified using the MinElute PCR Purification Kit (Qiagen). 1 µg of purified gap gene CDS served as template for dsRNA synthesis using the MEGAscript T7 Kit (Ambion). A gap gene dsRNA was diluted with injection buffer to a final concentration of 2 µg/µl and injected into female beetles.

## Notes

### Competing Interest Statement

The authors have declared no competing interest.

### Funding

- Deutsche Forschungsgemeinschaft

